# Chronic lower limb pain is not associated with a loss of inhibitory neurons in the human lumbar spinal dorsal horn

**DOI:** 10.64898/2026.09.15.751829

**Authors:** Olivia C Davis, Jane M Brandon, Keerthana Natarajan, Lucy He, Zaki Khan, Nethra Selvakumaran, Muhammad Babar, Aisha Alfayadh, Stephanie Shiers, Muhammad Saad Yousuf, Erin Vines, Peter Horton, Anna Cervantes, Tariq Khan, Geoffrey Funk, Andrew J Todd, Gregory Dussor, Theodore Price

## Abstract

The spinal dorsal horn is the primary processing site of nociceptive sensory input from the periphery. Excitatory spinal interneurons releasing glutamate can amplify this information before it is sent to the brain, whereas inhibitory neurons releasing GABA and/or glycine can suppress the outflow of nociceptive signals. An imbalance favoring excitation is thought to underlie certain aspects of chronic pain. Although rodent studies have identified spinal mechanisms underlying hyperalgesia and allodynia, little is known about the anatomical changes associated with chronic pain in the human spinal cord, a gap in knowledge we sought to address in this study.

Using immunohistochemistry and *in situ* hybridization on lumbar spinal cord tissue recovered from organ donors, we characterized neuronal size and density across the human dorsal horn and confirmed the presence of the human equivalent of the lateral spinal nucleus in many individuals. Chronic lower limb pain was not associated with changes in neuronal density in the dorsal horn. Likewise, the ratio of excitatory (*SLC17A6+*) to inhibitory (*PAX2+*) neurons remained consistent across laminae for age, sex and chronic pain state, providing no evidence for selective loss of inhibitory neurons with chronic pain in humans. We found no differences in the size or density of the postsynaptic markers Homer1 and gephyrin between groups, suggesting glutamatergic and GABAergic postsynaptic sites remain structurally stable. These findings provide a thorough evaluation of cellular anatomy of the human dorsal horn and form a foundation for future studies investigating neuronal changes that may contribute to chronic pain in humans.

**Summary:** The total number of neurons and the ratio of inhibitory to excitatory interneurons in the human spinal dorsal horn are not affected by chronic lower limb pain.

## Introduction

The dorsal horn (DH) of the spinal cord is a major processing site of nociceptive information arriving in the central nervous system from the periphery. Excitatory spinal interneurons release glutamate to amplify these signals, whilst inhibitory spinal interneurons release GABA and/or glycine to suppress this input. The balance of spinal excitation to inhibition is critical to maintain sensory thresholds, as an imbalance in favor of excitation drives chronic pain [2; 35]. Increased peripheral drive consequent to injury or inflammation can lead to enhanced glutamatergic signaling and synaptic strengthening in the DH, resulting in increased excitation of postsynaptic output neurons [18; 38; 57]. In contrast, loss of inhibitory tone can also favor excitation [8; 9; 34; 36]. Peripheral nerve axotomy has been proposed to lead to the loss of inhibitory DH interneurons in rodents [62], although this view was challenged by several stereological studies that showed no loss of inhibitory interneurons after peripheral nerve injury [31–33]. Whether inhibitory neurons are selectively lost in the DH of patients with chronic pain remains unknown.

Chloride gradient dysregulation has also been shown to contribute to spinal disinhibition. Peripheral nerve injury leads to a reduced expression of the potassium chloride cotransporter 2 (KCC2) in spinal neurons, altering the postsynaptic events following GABA_A_ receptor activation, by producing a reduced inhibitory effect compared to non-injured controls [9]. We recently showed a decrease in membrane-associated KCC2 immunofluorescence in the superficial DH of tissues from organ donors with chronic lower limb pain, showing conservation between species [10]. A decrease in substance P immunoreactivity has been observed in the ipsilateral DH following leg amputation [16], potentially due to the degeneration of dorsal root ganglia (DRG) neurons, which has been detailed in donors with diabetes-induced peripheral neuropathy (DPN) [43]. However, no other studies have investigated changes in spinal neuronal populations associated with chronic pain.

Homer1 and gephyrin are scaffolding proteins that form a vital component of glutamatergic and GABAergic/glycinergic postsynaptic densities, respectively [45; 53]. We have previously shown that the size of Homer1+ and gephyrin+ puncta is consistent across the human DH, indicating that these synapses are stable in normal conditions [11]. Homer1+ puncta have been shown to increase in size following focal two-photon glutamate uncaging, alongside dendritic spine increase [21], suggesting activity-dependent synaptic plasticity is associated with an enlargement of the postsynaptic density, reflected by increased Homer1+ punctum size. Synaptic alterations in the DH could also contribute to the maintenance of a chronic pain state, but this has never been assessed in humans.

Here we investigated neuronal structural and distribution differences in human spinal DH tissue from adults with no history of persistent pain compared to those with documented chronic bilateral lower limb pain. Our results find no evidence for the selective loss of inhibitory neurons and no changes in the size of glutamatergic or GABAergic synapses associated with chronic pain. While we acknowledge some limitations to the findings described here, our data guides future efforts to investigate other mechanisms that may contribute to chronic pain in humans.

## Methods

### Tissue procurement

All human tissue recovery procedures and ethical regulations were approved by the Institutional Review Board at the University of Texas at Dallas under protocol Legacy-MR-15-237. Spinal cords were procured from organ donors through a collaboration with the Southwest Transplant Alliance, an organ procurement organization (OPO) in Texas. The Southwest Transplant Alliance obtains informed consent for research tissue donation from first-person consent (driver’s license or legally binding document) or from the donor’s legal next of kin. Policies for donor screening and consent are those established by the United Network for Organ Sharing (UNOS). OPOs follow the standards and procedures established by the US Centers for Disease Control (CDC) and are inspected biannually by the Department of Health and Human Services (DHHS). The distribution of donor medical information follows HIPAA regulations to protect donor privacy. Spinal cords from the upper thoracic to sacral segments were surgically dissected from human organ donors within 1.5 - 3.5 hours of cross clamp and immediately frozen using crushed dry ice, as previously described [44]. Tissue blocks were stored at -80°C until use.

### Immunohistochemistry processing

The lower lumbar region (L4/L5) of spinal cords were dissected from the rest of the block using a sharp razor blade and embedded in sequential thin layers of OCT, ensuring the tissue remained fully frozen. Tissues from over 120 individual organ donors were cut into 20μm transverse sections using a Leica cryostat and mounted onto charged slides, then fixed in freshly depolymerized 4% paraformaldehyde for 20 minutes. Following this, sections were incubated in sequential concentrations of ethanol dissolved in distilled water (50%, 70%, 100%, and 100%) for 5 minutes each. Sections were allowed to air dry, then a hydrophobic barrier was drawn with an Immedge pen. Once the barrier was dry, sections were rehydrated with phosphate-buffered saline (PBS) and rinsed twice for 10 minutes with fresh PBS. All sections were then incubated with mouse IgG1 anti-NeuN (1:1000; Sigma-Aldrich, Cat. # MAB377; RRID: AB2298772) and left overnight in a dark moisture-controlled tray at 4°C. The following day slides were rinsed with PBS 3 times for ten minutes each, then reincubated with a goat anti-mouse IgG1 secondary antibody conjugated to Alexa Fluor 647 (1:500; Invitrogen; Cat. # A-21240) overnight in the dark at 4°C. After three final PBS rinses, slides were coverslipped using Prolong Gold (Invitrogen, Cat. # P36930) and kept in the dark at -20°C until scanning. Slides were scanned using an Olympus VS200 slide scanner with a dry x20 lens, and the tissue quality was assessed to allow the choice of high-quality tissue that could be used for analysis. High tissue quality was defined as having clear NeuN-immunolabelling throughout the spinal cord and no major holes or tissue damage to the DHs of the spinal cord resulting from surgical dissection, immunohistochemical processing, or ice crystal formation during the freezing process. Tissue sections were assessed for quality with the experimenter blind to age, sex and pain state.

From those with high-quality tissue, 18 age- and sex-matched donors were selected for further analysis (10 female, 8 male; 33 - 75 years of age; 10 non-pain and 8 with a clear medical history of lower limb pain; see Table 1 for donor demographic information). Fresh 20µm lower lumbar (L4/L5) sections were cut from tissue from each donor such that each slide contained multiple sections, which were more than 100µm apart in the rostrocaudal axis. Slides were processed for immunohistochemistry as described above. For cell cross sectional area analysis, new sections were incubated in mouse IgG1 anti-NeuN as above, whereas slides were incubated in guinea pig anti-Homer1 (1:500; Synaptic Systems; Cat. # 160 004, RRID: AB_10549720) and mouse IgG1 anti-gephyrin (1:500; Synaptic Systems; Cat. # 147 111; RRID: AB_2232546) for postsynaptic density measurements. An anti-guinea pig secondary antibody conjugated to Alexafluor 488 and an anti-mouse IgG1 secondary antibody conjugated to Alexafluor 647 were used to detect Homer and gephyrin-immunolabelling, respectively (1:500; Invitrogen; Cat. #A-11073 & A-21240). Secondary antibody controls were reacted simultaneously following the same protocol but excluding primary antibodies.

**Table 1:** Demographics of human organ donors used in this study. Parentheses in the Pain or Non-Pain column are Nageotte nodule scores from the DRG of the same donor from Shiers et al. 2025.

| Age (years) | Sex | Ethnicity | Cause of Death | PMI (hours) | Medical history of note | Pain or Non-pain |
| --- | --- | --- | --- | --- | --- | --- |
| 33 | M | White | Head trauma / MVA | 1.5 | Nothing of note. | Non-pain |
| 37 | F | Black | Head trauma / GSW | 3 | Nothing of note. | Non-pain |
| 41 | M | Black | Head trauma / GSW | 1.5 | Nothing of note. | Non-pain |
| 41 | F | White | Head trauma / MVA | 3 | Nothing of note. | Non-pain |
| 53 | F | White | Cerebrovascular accident / stroke | 3 | Nothing of note. | Non-pain |
| 54 | M | White | Anoxia / cerebrovascular accident | 2 | Nothing of note. | Non-pain |
| 64 | M | White | Cerebrovascular accident / stroke | 2.5 | Nothing of note. | Non-pain |
| 65 | F | Black | Cerebrovascular accident / stroke | 2 | Nothing of note. | Non-pain |
| 75 | M | White | Cerebrovascular accident / stroke | 2 | Nothing of note. | Non-pain |
| 75 | F | White | Cerebrovascular accident / stroke | 2 | Nothing of note. | Non-pain |
| 40 | F | Black | Anoxia / stroke | 2 | Rheumatoid arthritis of knees with pain. | Pain |
| 48 | F | Hispanic | Cerebrovascular accident / stroke | 2 | Diabetic peripheral neuropathy with lower limb pain and partial left foot amputation. | Pain (3) |
| 48 | F | White | Anoxia / cerebrovascular accident | 3.5 | Rheumatoid arthritis of knees with pain. | Pain |
| 52 | F | Black | Cerebrovascular accident / stroke | 3 | Diabetic peripheral neuropathy with lower limb pain. | Pain (4) |
| 53 | F | White | Anoxia / cerebrovascular accident | 2 | Fibromyalgia with widespread pain. | Pain |
| 60 | M | White | Anoxia / cerebrovascular accident | 1.5 | Peripheral neuropathy of the legs with chronic pain. | Pain (4) |
| 60 | M | Black | Anoxia / cerebrovascular accident | 2 | Diabetic peripheral neuropathy with lower limb pain. | Pain (4) |
| 63 | F | White | Anoxia / stroke | 2 | Osteoarthritis of lower limbs with pain. | Pain |

### Cell cross-sectional area analysis and identification of laminar boundaries

For cell cross-sectional area analysis, at least three sections through the entire DH of each donor were scanned with a dry x20 lens on an Olympus VS200 slide scanner microscope. A blank channel (∼560nm) with no immunolabelling was also scanned to view the autofluorescent pigment lipofuscin. Scanning settings were kept the same between all sections and donors. Secondary antibody controls were scanned using the same settings but were not analyzed. Images were then opened with QuPath–0.6.0 software and the grey matter of the dorsal horn was outlined using the soma and neuropil NeuN-immunolabelling, together with the autofluorescent signal and background myelin staining. Within this region, every NeuN-immunoreactive (NeuN-IR) cell body, in which the nucleolus was visible (seen as a small region of little to no NeuN-immunoreactivity within the nucleus of a NeuN-IR cell), was selected and then the outline of the NeuN-immunostaining was manually traced to create a polygon for each neuron using the brush tool. Once all NeuN-IR cell bodies were traced, Rexed laminar boundaries [37] were drawn using the density and morphology of these neurons following previously published anatomical landmarks and guides for the human spinal cord [14; 39; 50]. Briefly, lamina II was defined as the dense band of cells across the entire width of the dorsal horn, and cells in this region often had more contrast against a darker background as there was less autofluorescence from myelin in this lamina (the substantia gelatinosa). Lamina I was the region between lamina II and the dorsal edge of the white matter. For analysis, any rare neurons in the white matter dorsal to the lamina I boundary and those in Lissauer’s tract were included in lamina I. Lamina III was directly ventral to lamina II and contained slightly larger cells that were less densely packed than those in lamina II. The laminae III/IV boundary could be seen as a change in neuronal density, with lamina IV being less dense again and containing several very large neurons. The ventral edge of the lamina IV boundary was drawn at a further change in NeuN-IR neuropil and cell density, as lamina V often had less background neuropil immunolabelling and dense clusters of small cells could also be seen, which were largely absent in lamina IV. The laminae V/VI boundary is impossible to define in human spinal cord [39; 50], so these laminae were analyzed together, and the ventral edge of lamina VI was drawn at a further change in neuron density observed around the base of the dorsal horn. The reticulated lamina V (RLV) region was drawn at the lateral edge of laminae V/VI where bundles of myelinated fibers were visible in the background staining. Finally, several neurons were found in the lateral white matter. While the lateral spinal nucleus (LSN) has been depicted anatomically on a human spinal cord atlas [55], the existence of the human equivalent of the LSN has not been substantiated by direct evidence of a discrete neuronal population in the corresponding region, nor has its anatomical organization been systematically characterized across multiple individuals. As the number and pattern of NeuN-IR cells in the white matter varied between donors we combined all neurons in this area in a region termed dorsolateral white matter (DLWM) for comparison. The area of each laminar boundary as well as the cross-sectional area of all traced neurons within each region were exported to a csv file. Numbers were combined and averaged from multiple sections through the dorsal horn for each donor.

The entire spinal cord sections were also imaged at a x20 magnification using the Olympus VS200 slide scanner microscope. These images were opened in Qupath–0.6.0 software and the entire outline of the tissue sections was traced to provide the cross-sectional area of the spinal cord for each donor. The area of each DH was also normalized by dividing by the cross-sectional area of the whole spinal cord, to calculate the relative proportion of the DH as a part of the whole for each section analyzed.

### Postsynaptic protein diameter analysis

Measurements of postsynaptic density proteins required high magnification images and so this tissue was scanned using an Olympus FV4000RS microscope with a x60 oil-immersion lens and a 2.28x zoom (optimal for the aperture and lens used) to reach a resolution of 0.09µm per pixel. The laser settings and scanning conditions were kept the same between all sections and donors. Three scans (one medial, one central and one lateral to account for innervation differences) of approximately 100µm^2^ were taken from laminae I, II and deep dorsal horn on both sides of the spinal cord section, totaling 9 scans per section for multiple sections per donor. Each scan was a z-series throughout the tissue thickness (20µm), with a z-interval of 0.3µm. A blank channel (∼560nm) was also scanned along with the channels containing Homer1 and gephyrin immunolabelling to reveal autofluorescence. Lamina II was identified due to the visibility of the substantia gelatinosa under the confocal eye piece (not scanned) and dense lipofluorescent signal seen from the high neuronal density across the width of the grey matter. Lamina I was defined as the region dorsal to the substantia gelatinosa up to the edge of the white matter. The deep dorsal horn (laminae VI/V) region was selected as that sits approximately 500µm dorsal to the central canal, where lipofuscin signal from large neurons was visible. Secondary antibody control sections were scanned using the same settings but were not analyzed. For analysis of glutamatergic postsynaptic densities, scans were opened in ImageJ (v1.54) and viewed with a 10µm squared grid overlaid. The channels containing lipofuscin and Homer1-immunolabelling were initially viewed, then 50 Homer1-IR puncta (lacking lipofuscin, and therefore not autofluorescence) that contacted the gridlines were selected. Each of these selected profiles was then assessed in the z-axis and measured at the widest point in diameter using the line tool. Following this, the same analysis was conducted with the channels for lipofuscin and gephyrin-immunolabelling visible to assess inhibitory synapse sizes. Numbers were combined from multiple dorsal horns for each donor.

### Fluorescent in situ hybridization processing and analysis

In initial tests, we trialed numerous antibodies raised against fast neurotransmitters or the transcription factors PAX2 and LMX1B, which can be used to identify inhibitory and excitatory neurons (see Supplemental Table 1 for details), but found that these did not give consistent labelling of spinal neurons in human spinal cord sections. Therefore, we used *in situ* hybridization to label inhibitory and excitatory neuronal somas. Transverse spinal cord sections (20µm) were cut and mounted onto charged slides as above. Multiple sections more than 100µm apart in the rostrocaudal axis were placed onto the same slide for each donor and left overnight at -80°C to aid tissue adherence. In situ hybridization was performed with a RNAscope fluorescent multiplex reagent kit 320850 (ACD BioTechne; Cat. # 323100) following manufacturer instructions. To visualize inhibitory and excitatory neurons, probes for *PAX2* (Cat. #P36930) and *SLC17A6* (Cat. # 415671-C3) were used, respectively. After RNAscope processing, slides were incubated overnight with mouse anti-NeuN (1:500) as described above. Slides were then rinsed three times with PBS and reincubated with a goat anti-mouse IgG1 secondary antibody conjugated to Alexa Fluor 488 (1:500; Invitrogen; Cat. # A-21121). The following day, sections were incubated with DAPI (1:2000; Cayman Chemicals; Cat. # 14285) for 5 minutes then rinsed twice and mounted using Prolong Gold (Invitrogen; Cat. # P36930). Slides were allowed to cure for 24 hours in the dark before scanning. Positive and negative control probes were also tested on other sections at the same time.

Sections were scanned through the full thickness with a 0.91 μm z-step on an Olympus FV4000 confocal microscope using a x20 dry lens with a 2x zoom. Multiple scans were taken from laminae I, II and the deep dorsal horn (the laminae IV/V boundary) from multiple tissue sections. Laminar regions were determined based on NeuN-immunolabelling as described above. Z-series of images were then opened using Neurolucida software (MBF Bioscience, VT, USA). With only the channels containing NeuN-immunolabelling and DAPI visible, NeuN-IR cell bodies were selected before the remaining channels were visualized. Each NeuN-IR cell was assessed for the presence or absence of *PAX2+* and *SLC17A6*+ puncta. The autofluorescent pigment lipofuscin could be visualized as a perfect overlap of signal in the channels containing *PAX2* and *SLC17A6* probes and NeuN-immunolabelling, as lipofuscin fluoresces throughout the spectrum captured [5; 22]. All three channels were therefore visualized at the same time to delineate true probe signal from autofluorescent artefact. These markers of excitatory and inhibitory neurons are highly expressed such that neurons often contained either >15 transcripts or no transcripts at all of *PAX2* or *SLC17A6*. A cell was therefore defined as being positive if 3 or more *PAX2*+ or *SLC17A6*+ puncta were present. Each neuron was assessed in the Z-axis to ensure accurate separation between cells in close contact. Numbers were combined from multiple dorsal horns for each donor.

### Western Blot

Lumbar spinal cord tissues from the same 18 donors (see Table 1 for donor demographics) were stored at −80°C until they were processed. Approximately 1mm transverse sections were rapidly dissected from the lower lumbar region of spinal cord, rostral to the tissue sections used for immunohistochemistry and in situ hybridization, with a fresh razor blade. The ventral half of the block (below the central canal) was removed, leaving the grey matter of the dorsal horns and the surrounding white matter. These dorsal segments (∼100mg) were processed for western blots as previously described [12]. Briefly, tissues were manually minced with scissors in ice-cold lysis buffer consisting of RIPA buffer (Thermofisher Scientific; Cat. # 89901) with a phosphatase inhibitor cocktail added immediately before use (Sigma Aldrich; Cat. # P8340, #P5726, #P0044; total volume of lysis buffer per sample: 750μL). The tissue and lysis buffer mix was further crushed in homogenizer glass tubes for several minutes on ice. Finally, samples were centrifuged at 10,621 g for 10 min at 4°C. Protein concentration was quantified by the Bicinchoninic acid (BCA) assay (ThermoFisher Scientific; Cat. #23225). The samples were aliquoted and denatured at 95°C for 5 min after the addition of 4x Laemmli with β-mercaptoethanol (Bio-Rad; Cat. # 1610747) and kept at -80 °C until use.

A total of 30 µg of protein was resolved by denaturing in 10% and 15% sodium dodecyl sulfate-polyacrylamide gel electrophoresis and were then transferred overnight to polyvinylidene difluoride (PVDF) membranes (Millipore Sigma; Cat. # IPFL00010). The PDVF membranes were blocked for 2 hours at room temperature in 5% bovine serum albumin (BSA) in 1X Tris buffer with Tween20 (TBST; 150 mM NaCl, 200 mM Tris at pH 7.4, 0.1% Tween 20). They were then incubated overnight at 4°C in either guinea pig anti-Homer1 (1:500; Synaptic Systems; Cat. # 160 004; RRID: AB_10549720), mouse IgG1 anti-Gephyrin (1:500; Synaptic Systems; Cat. # 147 111; RRID: AB_887719), rabbit anti-GABA_A_ α2 (1:500; Synaptic Systems; Cat. # 224 103; RRID: AB_2108839) or rabbit anti-GAPDH (1:1000; Cell Signaling; Cat. # #2118; RRID: B_561053) diluted in TBST + 5% BSA. Membranes were rinsed three times with TBST + 5% BSA for 10 minutes at room temperature, then incubated in secondary antibodies for 2 hours at room temperature. Secondary antibodies were also diluted in TBST + 5% BSA (1:10,000) and were either goat-anti-rabbit IgG (Jackson Immunoresearch Laboratories; Cat. # 111-035-003), goat anti-guinea pig (Jackson Immunoresearch Laboratories, Cat. #106-035-003; RRID: AB_2337402) or goat anti-mouse IgG (Jackson Immunoresearch Laboratories, Cat. #115-035-003; RRID: AB_10015289) secondary antibody conjugated to horseradish peroxidase. Membranes were rinsed a final three times with TBST + 5% BSA then processed for protein signal detection using the enhanced chemiluminescence system (ECL plus; Thermo Scientific; Cat. # 32132). Bands were quantified using Image Lab Software Version 5.2.1 (Bio-Rad). Following imaging for each primary antibody, the blots were rinsed with TBST then incubated in stripping buffer (Thermofisher Scientific; Cat. # 46430X4) and shaken lightly to ensure even distribution of buffer for 21 minutes at 30°C. Membranes were rinsed 3 more times with TBST then reacted for the next antibody in the same way described.

Glyceraldehyde-3-phosphate dehydrogenase (GAPDH) is a constitutively expressed housekeeping gene, used previously as a control protein for western blots examining protein expression in the spinal cord after peripheral nerve injury in rodents [46]. We compared GAPDH chemiluminescence bands (∼37kDa) between our non-pain and pain samples and found there was no significant difference (unpaired t-test, p = 0.8), therefore chemiluminescence of Homer1 (45kDa), Gephyrin (93kDa) and GABA_A_ α2 (50kDa) were normalized to the GAPDH bands for each sample.

### Statistical analysis

All statistical analysis was conducted using GraphPad Prism version 10.0.2 (GraphPad Software, San Diego, CA, USA). Values were averaged for each donor prior to statistical analysis, such that each donor represented a single biological replicate. Comparisons between two independent groups (sex, age, or pain state) were performed using unpaired t-tests. For comparisons between pain and non-pain donors within individual laminae, multiple t-tests with Holm–Šídák correction for multiple comparisons were used. To assess postsynaptic marker size across laminae and pain states, a 2way-ANOVA with Tukey’s multiple comparison test was used. P values of less than 0.05 were considered significant. All data are shown as mean ± standard deviation.

## Results

### Neuronal size and density across the spinal dorsal horn

High-quality tissues from 18 age-matched donors of both sexes were selected from more than 120 organ donors that were initially assessed. Donors were selected following blinded assessment of tissue quality to form an age- and sex-matched cohort where possible. From these 18 donors, 10 had no history of persistent pain (non-pain; 5 females and 5 males; 33 – 75 years old) and 8 donors had documented chronic bilateral lower limb pain (pain; 6 females and 2 males; 40 – 63 years old; see Table 1 for donor demographic information). Following immunohistochemistry for NeuN, the background staining in the white matter of the spinal cord was variable between donors; however, the quality of NeuN immunostaining was consistent when viewed at a single-cell resolution, clearly staining the cell soma of neurons across the tissue section (see Figure 1 and Supplemental Figures 1-3 for representative high and low magnification images from each donor). The cross-sectional area of spinal cord tissue sections did not differ between non-pain donors when grouped by sex or age (see Supplemental Table 2 for detailed summary of results and statistical analyses). For example, there was no difference in spinal cord cross-sectional area when comparing under 60 (n = 6; 3 females, 3 males; 33 – 54 years) to over 60 years of age (n = 4; 2 females, 2 males; 64 – 75 years). Likewise, there was no difference in the cross-sectional area between non-pain donors and those with chronic lower limb pain (Supplemental Table 2).

**Figure 1:**
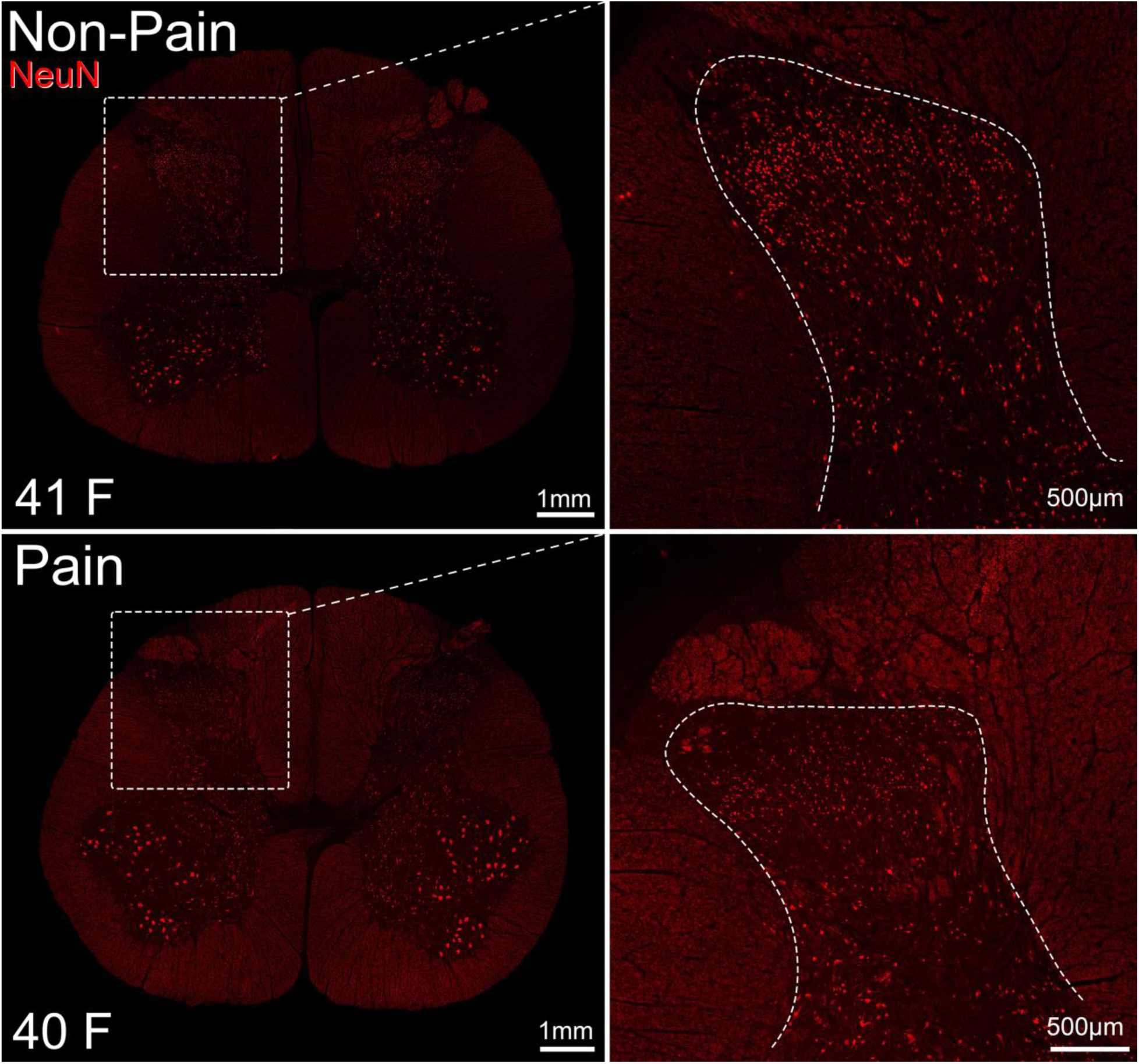
Representative NeuN immunolabelling in the dorsal horn of transverse lower lumbar spinal cord sections from organ donors with and without chronic lower limb pain. Images are a single optical section; insets are cropped from the images on the left. x20 magnification. F = female. Numbers equate to age in years.

While there was individual variation in the shape of the grey matter of the spinal DH (Figure 1, Supplemental Figures 1 & 2), there was no significant difference in the absolute DH area between sexes or different ages in non-pain donors, or between those with and without pain (Figure 2A). We then calculated the absolute area of the DH as a proportion of the cross-sectional area of the entire spinal cord. Although the absolute DH area did not differ, this relative DH area was significantly smaller in donors with chronic pain, however, this effect was modest with an average of 0.069 ± 0.0069 in non-pain and 0.062 ± 0.0049 in pain donors (p = 0.02; Figure 2B). This is unlikely due to differences in the size of the grey matter between the L4 and L5 spinal cord segments, as comparable proportions of tissue sections from non-pain and pain donors were from each region (80% of non-pain donors and 75% of pain donors had tissue sections from the L5 region, while the rest were from the L4 segment). The number of neurons across the DH also varied between individuals but did not significantly differ between sexes, different ages, or pain states (Figure 2C), meaning the packing density of neurons across the DH was consistent between groups (Figure 2D). The average size of neuronal cell bodies was comparable between all groups (Figure 2E & F), with an average neuronal soma area of 183.2 ± 172.0µm^2^ in non-pain donors and 169.4 ± 170.9µm^2^ in those with chronic lower limb pain. Again, when normalized to the cross-sectional area of the spinal cord, there were no differences in the average relative neuron soma size between groups.

**Figure 2:**
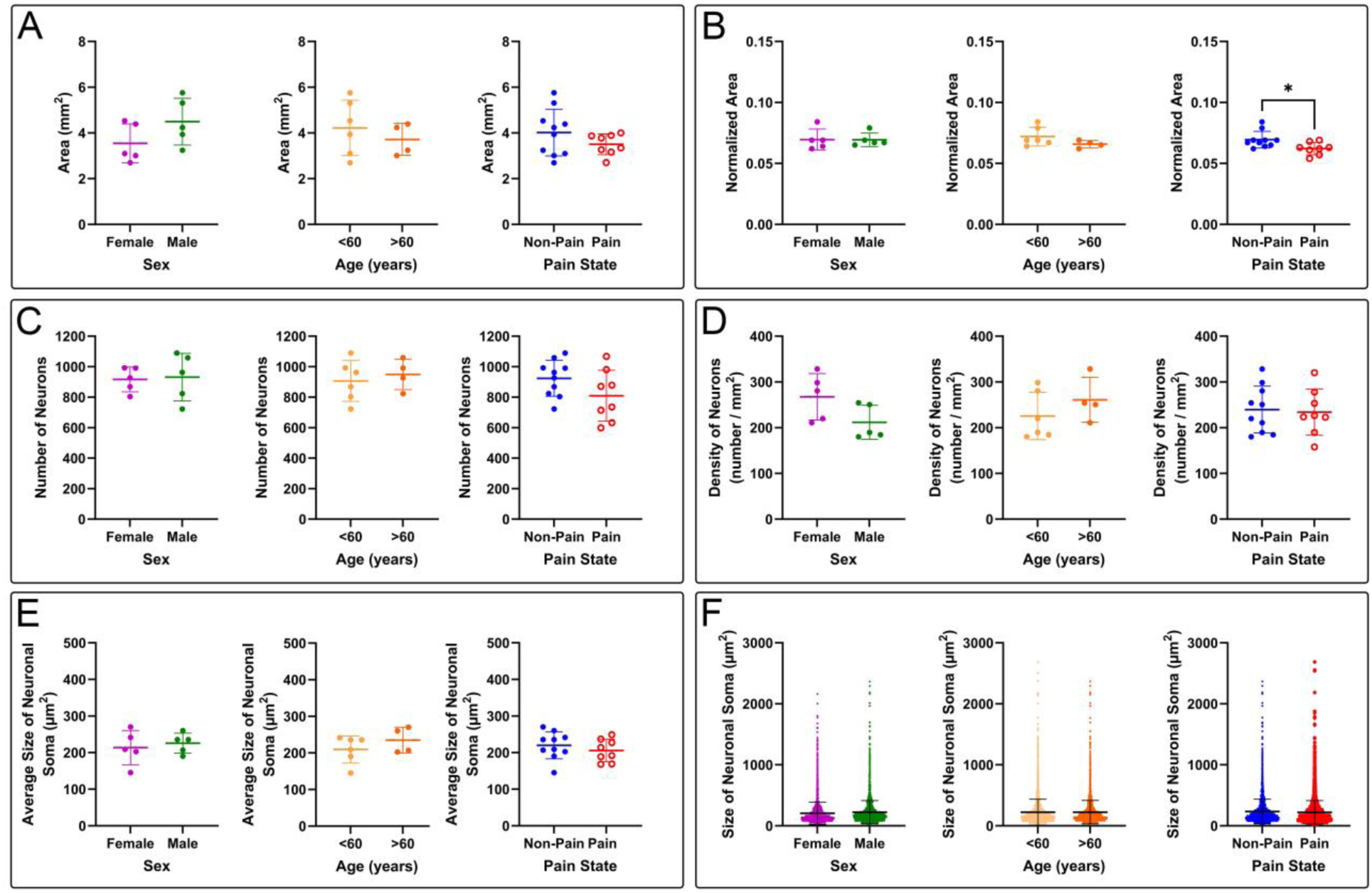
The size and neuronal packing density of the spinal dorsal horn are not affected by sex, age or chronic lower limb pain. The cross-sectional area of the grey matter of the spinal dorsal horn (A), the relative dorsal horn area normalized to the cross-sectional area of the spinal cord (B), the number of NeuN-immunolabelled neurons (C), the packing density of neurons (D), the average and range of the neuronal soma sizes (E & F) are comparable between donors with no medical history of chronic pain (non-pain) when grouped by sex or age. Most of these parameters are affected not by chronic lower limb pain (pain), however the normalized area of the spinal dorsal horn is significantly smaller in pain donors (p = 0.02). P > 0.05 for all other comparisons.

In the rodent, projection neurons are significantly outnumbered by local interneurons, accounting for 1% of all spinal neurons [7]. Despite their rarity, projection neurons can be identified in the SDH as they have been shown to have significantly larger cell bodies than interneurons in both mice and rats [1; 6]. We noted the presence of very large, rare cells in lamina I and the deep dorsal horn of all tissue sections analyzed. The largest 1% of human SDH neurons measured were found to be 885.6µm^2^ - 4737.7µm^2^ (62,041 neurons in total), providing a size threshold for identifying candidate projection neurons in future analyses. In contrast, the vast majority of neurons (∼90%) had a soma cross-sectional area of less than 300µm^2^.

### Neuronal size and density across different dorsal horn laminae

Rexed laminar boundaries were determined for each donor following previously defined guides for the human spinal cord and using neuronal soma morphology and packing density as shown in Figure 3 [14; 37; 39; 50]. Despite differences in DH shape between individual donors, the proportion of each laminar region within the DH did not differ with age, sex or pain state. Further, none of the anatomical measurements taken including laminar region area (Figure 4A-B), number of neurons (Figure 4C), packing density (Figure 4D) and neuronal soma area (Figure 4E-4F) differed between any groups, in any laminar region (p > 0.05 for all pairwise comparisons, see Supplemental Table 2 for detailed results and statistical analyses).

**Figure 3:**
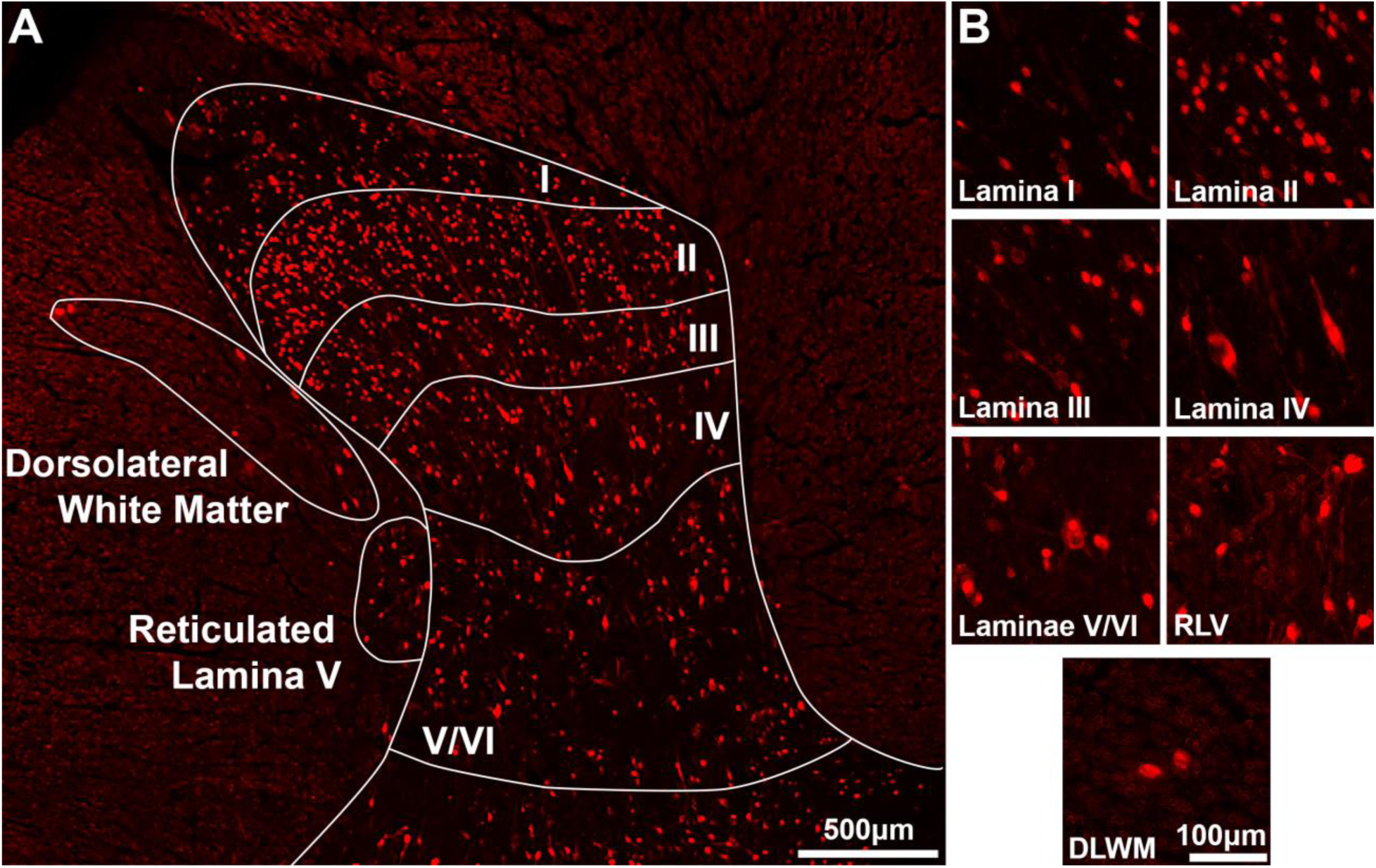
Laminar boundaries drawn on the human lower lumbar dorsal horn using the density and morphology of NeuN-immunoreactive cell bodies. Seven regions were annotated for further analysis based on NeuN immunolabelling (A). Briefly, lamina I contained both small and large cells at the edge of the white matter (B). Lamina II contained the smallest, roundest cells that were densely packed together across the dorsal horn while lamina III was slightly less dense and contained slightly larger cells. Lamina IV contained several very large cells and fewer smaller cells. laminae V and VI cannot be separated in human spinal cord so were grouped together and contained large cells as well as clusters of small, round cells that are largely absent in lamina IV. A region at the lateral edge of the deep dorsal horn with high background staining contained relatively few neurons (reticulated lamina V / RLV) and every donor analyzed had neurons in the white matter adjacent to the dorsal horn, presumably belonging to several ascending tracts and were grouped together (dorsolateral white matter / DLWM). Image is a single optical section; insets in B are taken from the figure in A.

**Figure 4:**
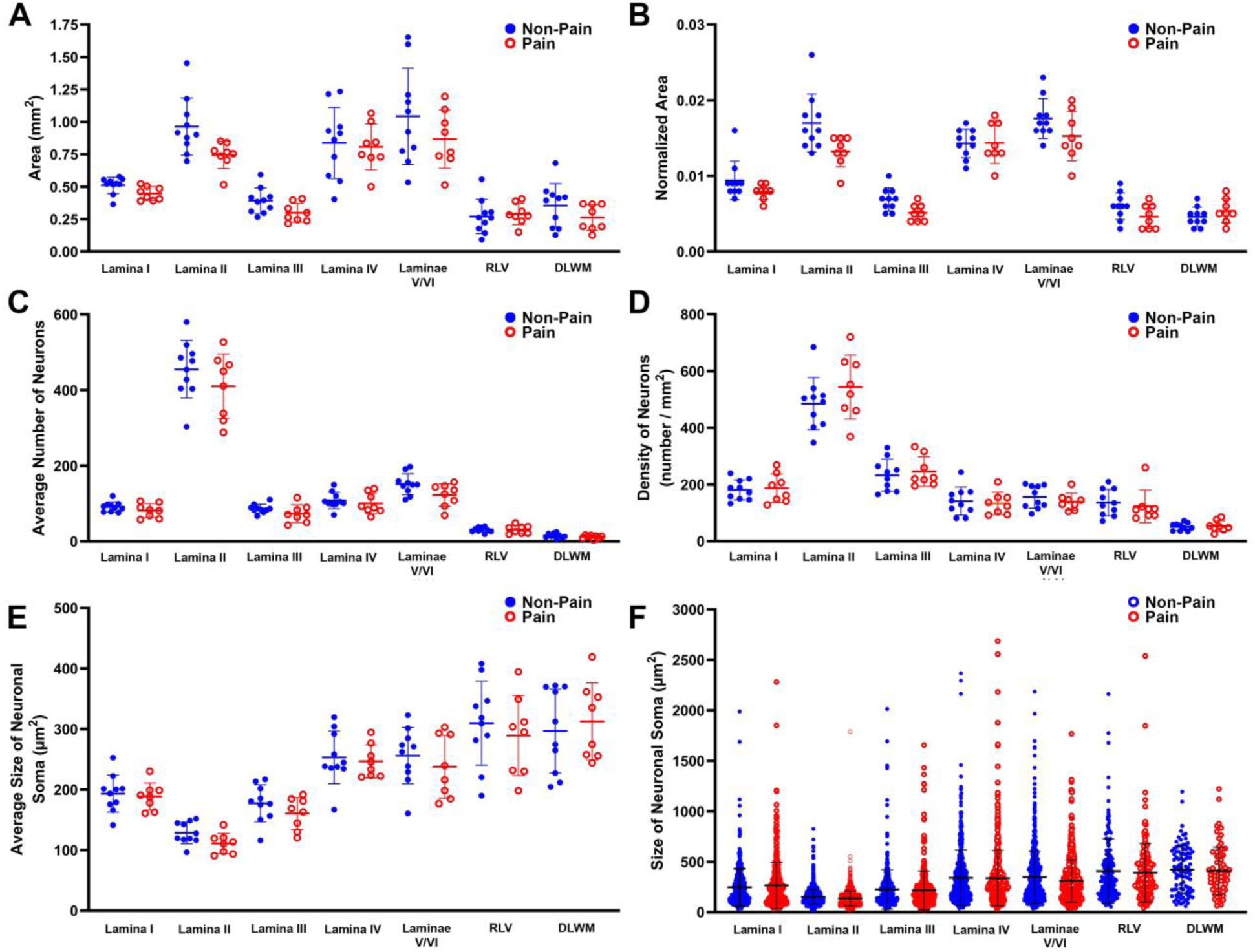
Neuronal density and size vary across laminar boundaries, but not between donors with and without a history of chronic lower limb pain. The cross-sectional area of each laminar boundary (A), the area normalized to the cross-sectional area of the spinal cord (B), the number of NeuN-immunolabelled neurons (C), the packing density of neurons in the dorsal horn (D) and the average neuronal soma size (E) are comparable for each region analyzed between donors with no medical history of chronic pain (non-pain) and those with documented chronic lower limb pain (pain). The spread of soma sizes varies across the dorsal horn (F) highlights the presence of very large cells in lamina I, the deep dorsal horn and the white matter adjacent to the dorsal horn. Adjusted P values > 0.05 for all pairwise comparisons between non-pain and pain for each region.

#### Lamina I

As shown in Figure 3, lamina I was defined as the region of grey matter between the white matter boundary and a dense plexus of cells that comprise lamina II. Lamina I contained mostly small neurons and occasional neurons with a very large soma area, ranging from 17.3 - 2550.6µm^2^ across all donors (Figure 4). Taking the potential projection neuron threshold calculated above of approximately 885µm^2^, 1.1% of neurons in lamina I fell within this category (67 / 6156 neurons).

#### Lamina II

Lamina II was defined as a dense band of small, spherical neurons with a fairly circular cross-section across the width of the grey matter (Figure 3). There was sometimes a greater contrast between NeuN-IR cells in this region compared to the background staining than in the rest of the dorsal horn, presumably due to a lack of myelin in this lamina (the substantia gelatinosa). Lamina II contained the highest number of neurons across the DH, corresponding to a packing density almost three times greater than that of any other laminar region (Figure 4B). Large neurons (>885µm^2^) were almost entirely absent from lamina II, accounting for less than 0.03% of all neurons measured in this region (8 / 30,144) and neurons in this region had the smallest average soma area within the entire DH (129.5 ± 64.0µm^2^ in non-pain donors, 112.4 ± 61.3µm^2^ in those with pain).

#### Lamina III

The lamina II / III border could be identified by a decrease in neuronal density and an increase in soma size, as lamina III contained slightly larger neurons that were also less densely packed than those in lamina II (Figure 3). The neurons in lamina III often had a more elliptical soma cross section, with the widest diameter occurring in the dorsoventral axis. In contrast to rodents, this boundary was rarely straight and instead occurred at an uneven distance from the laminae I/II border when viewed across the width of the grey matter. Among the first four laminae, lamina III had the smallest area, with a similar number of neurons to lamina I. Large neurons were more numerous in lamina III than lamina II, with 58 / 5682 neurons having a soma area greater than >885µm^2^, although these still only accounted for approximately 1% of neurons in this region.

#### Lamina IV

The dorsal boundary of lamina IV could be identified as a further decrease in neuronal density compared to lamina III, with many more large neurons and fewer smaller neurons (Figures 3 & 4). Large neurons accounted for 2.7% of lamina IV neurons (196 / 7324) and often had elongated cell bodies with dorsally directed dendrites labelled with the NeuN antibody. Due to this neuropil staining, lamina IV often contained a background of NeuN-immunolabelling, which was absent in lamina V and could be used to determine the ventral boundary of lamina IV.

#### Laminae V/VI

While laminae V/VI also contained many large neurons (2.1% of all lamina V neurons, 199 / 9698), there were frequent clusters of small neurons that were not present in lamina IV, where small cells were distributed without a discernable pattern. This neuronal organization pattern was used to draw the lamina IV/V boundary and overlapped with the cessation of the neuropil staining observed in lamina IV of some donors. Laminae V and VI cannot be distinguished in human lumbar spinal cord [39], so these were kept as one region. The ventral boundary of lamina VI could be seen as a slight curved gap in the neuronal density at the base of the dorsal horn. Lamina V/VI was the largest laminar region, although both lamina II and lamina IV individually covered similarly large areas of the DH on average (Supplemental Table 2).

#### Reticulated lamina V (RLV)

While the V/VI laminar boundary cannot be defined, white matter reticulations created a web-like pattern of autofluorescence in the lateral edge of laminae V and VI. This lateral reticulated lamina V (RLV) region has been shown to contain a specific population of projection neurons involved in sensory processing [3; 4; 20]. This region had the highest proportion of large cells, with 5.9% of neurons having a soma greater than >885µm^2^, although the actual number of large neurons was relatively low as there were only 2254 neurons in RLV measured from all 18 donors.

#### Dorsolateral white matter (DLWM)

Finally, all donors contained neurons that were not located in the grey matter of the dorsal horn but could be found instead in the adjacent lateral white matter. These cells often ran parallel to the edge of laminae III-IV, although neurons were also located lateral to lamina II in a region potentially analogous to the LSN in all donors. As there has been no formal quantitative stereological studies of the LSN in humans, and some neurons appeared between the LSN and RLV in an intermediate zone (see Figure 3), all neurons in the white matter were grouped in a region called the dorsolateral white matter (DLWM). Again, a high proportion of neurons (4.4%) in this region were considered large (>885µm^2^), although the number of neurons and therefore packing density in this region were the lowest of any measured.

### Ratio of inhibitory and excitatory dorsal horn neurons

Since the total number of neurons in the dorsal horn was unchanged, we next tested whether chronic pain was associated with alterations in the proportion of inhibitory and excitatory interneuron populations in the superficial and deep dorsal horn. Using fluorescent *in situ* hybridization, we found that just under 40% of neurons in lamina I of non-pain donors contained *PAX2*+ transcripts, but not *SLC17A6* and were therefore inhibitory neurons (Figure 5A-C; see Supplemental Table 3 for detailed results and statistical analyses). This was slightly lower in lamina II, with approximately one third of neurons being *PAX2*+/*SLC17A6*- (Figure 5D-E). The highest proportion of inhibitory neurons was identified in the deep dorsal horn at the laminae V/VI boundary with ∼47% of neurons containing *PAX2*+ puncta only (Figure 5F-G). The vast majority of the remaining neurons contained *SLC17A6*+ puncta and were therefore glutamatergic, although occasional neurons were found in all donors that contained both *PAX2*+ and *SLC17A6*+ transcripts, consistent with transcriptomic studies describing mixed molecular phenotypes in human spinal cord [13; 59; 61]. A small proportion of neurons were also found to contain neither *PAX2*+ or *SLC17A6*+ transcripts. The proportion of inhibitory (*PAX2*+/*SLC17A6*-), excitatory (*PAX2*- /*SLC1716*+), mixed (*PAX2*+/*SLC17A6*+) and neither *PAX2*+ or *SLC17A6*+ neurons were not significantly different in any laminar region when grouped by sex or age in non-pain donors (p > 0.05 for all pairwise comparisons, see Supplemental Table 3, Figure 5B-G).

**Figure 5:**
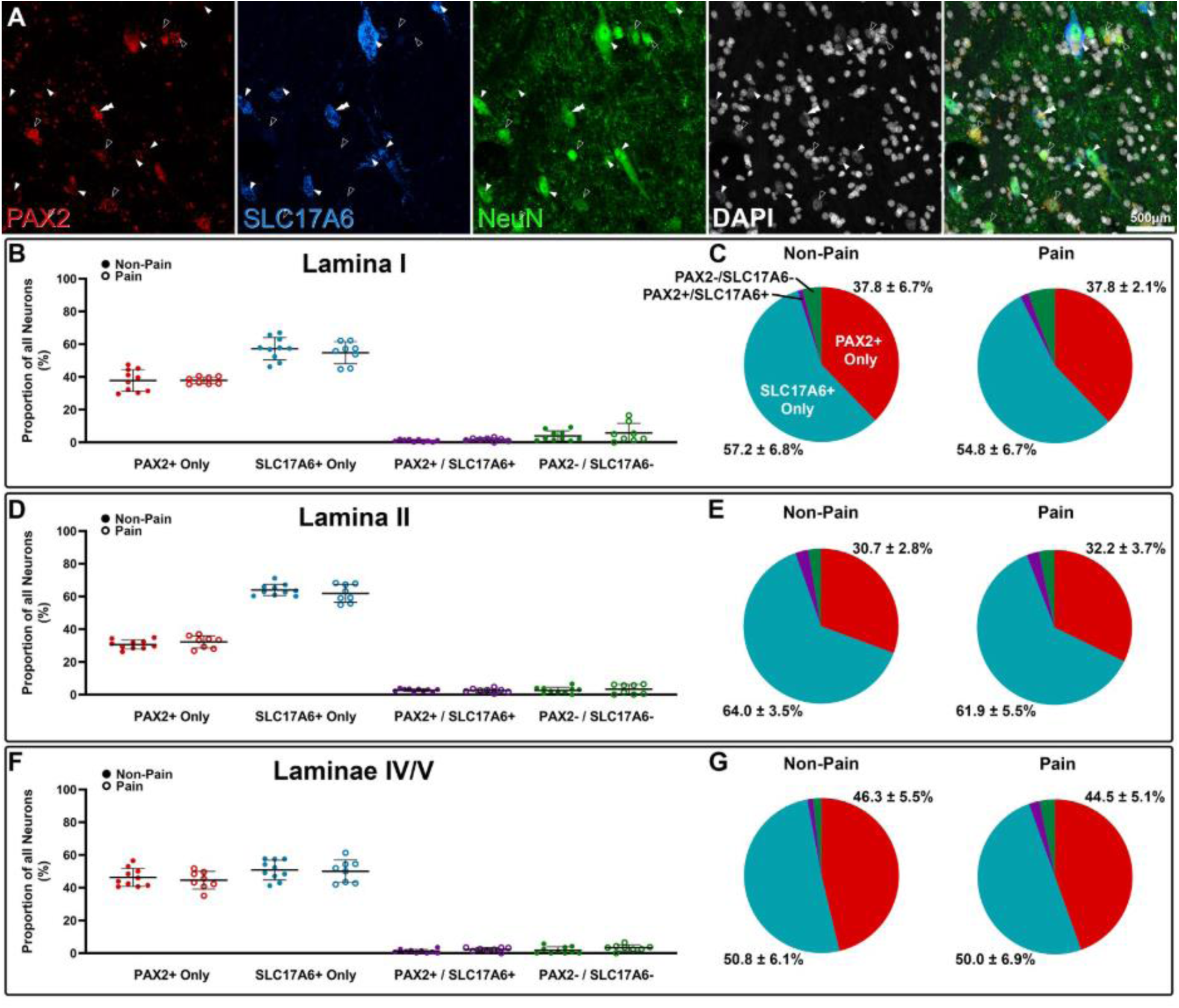
Chronic pain does not affect the ratio of inhibitory to excitatory neurons in the human dorsal horn. Representative image from the laminae IV/V boundary (A) showing fluorescent *in situ* hybridization with probes against *PAX2* (red) and *SLC17A6* (blue), in combination with NeuN-immunolabelling (green) and DAPI (white) to label inhibitory (empty arrowhead) and excitatory neurons (filled arrowhead), respectively. Occasionally, neurons contained transcripts for both *PAX2* and *SLC17A6* (double filled arrowheads). Approximately 38% of neurons in lamina I (B&C), just under a third of neurons in lamina II (D&E) and over 40% of neurons on the boundary of laminae IV and V (F&G) contain *PAX2*+ transcripts only and are therefore inhibitory. The vast majority of the remaining neurons in all regions were excitatory (*SLC17A6*+) and there was no difference in the proportion of inhibitory to excitatory cells between non-pain and chronic pain donors in any region (P > 0.05 for all comparisons). Image shown is a maximum projection of 7 optical sections at x20 magnification.

Similarly, the proportion of inhibitory and excitatory neurons was not significantly different in donors with chronic pain conditions in comparison to non-pain donors, in either lamina I, lamina II, or the deep dorsal horn. The proportion of neurons containing both *PAX2*+ and *SLC17A6*+ transcripts was also comparable between non-pain and pain donors in all regions (Figure 5), suggesting these mixed cells do not phenotypically switch to a single fast neurotransmitter following chronic pain. Together, these data provide no evidence for a change in the proportion of inhibitory and excitatory interneurons associated with chronic lower limb pain. Since total neuronal number did not differ between groups, unchanged proportions also indicate no detectable loss of inhibitory spinal neurons with chronic pain.

Out of 8 pain donors, 4 had pain induced by peripheral neuropathy (3/4 due to diabetes-induced peripheral neuropathy), 3 had pain resulting from arthritic conditions (1 rheumatoid, 2 osteoarthritis) and the remaining donor had fibromyalgia with widespread pain. The ratio of inhibitory to excitatory neurons in both the superficial and deep dorsal horn was comparable in donors with neuropathy in the lower limbs compared to those with arthritic medical histories (Supplemental Figure 4, Supplemental Table 3), although future studies should confirm this with greater numbers to make stronger conclusions.

### Synaptic protein architecture and density

The average size of Homer1-IR puncta in the superficial laminae was consistent between sexes and with age in non-pain donors (Figure 6A), consistent with previous reports [11]. Likewise, the average size of Homer1-IR puncta was found to be similar across all laminae and did not differ between pain and non-pain donors (Figure 6B; see Supplemental Table 4 for full results and statistical analyses; p > 0.05 for all comparisons). Gephyrin-IR puncta were a comparable size to Homer1-IR profiles (∼0.5µm) and showed no significant effect of pain (two-way ANOVA with Tukey’s multiple comparisons, p=0.22), with no pairwise differences between non-pain and pain in any laminar region (Figure 6C). Interestingly, gephyrin-IR puncta were significantly larger in the deeper dorsal horn than in the superficial laminae in both non-pain and pain donors (p value for pairwise comparisons <0.05, Figure 6C, Supplemental Table 4).

**Figure 6:**
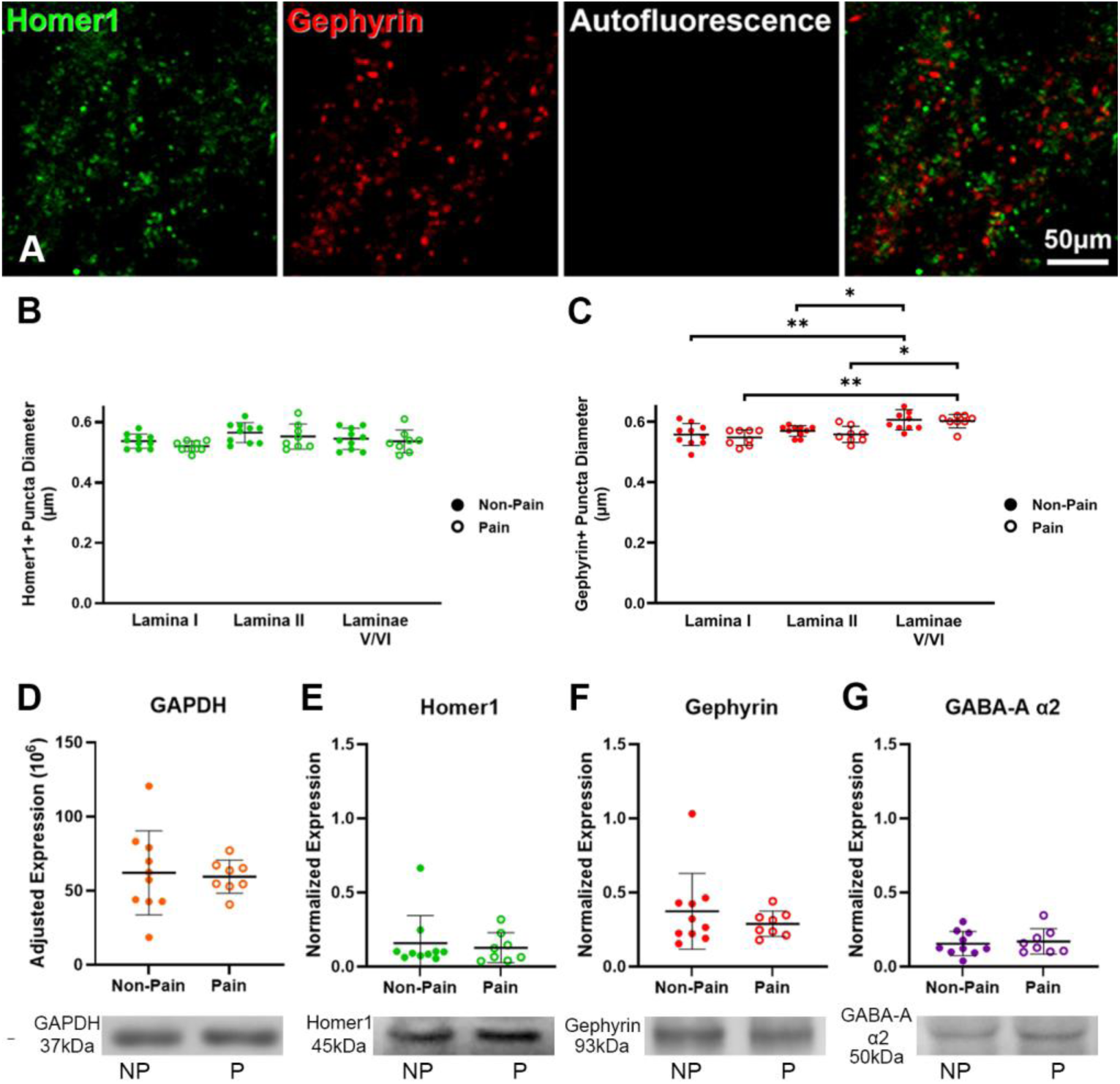
Chronic lower limb pain does not alter the size or quantity of either excitatory or inhibitory post synaptic density markers. Single optical image from lamina II showing Homer1 (green) and gephyrin (red) immunolabelling at x60 magnification that is not autofluorescence or artefact as shown in white (A). The size of Homer1-immunoreactive profiles is not altered with chronic lower limb pain in any region of the spinal cord (B). Gephyrin-immunoreactive puncta were larger in the deep dorsal horn compared to the superficial laminae in both pain and non-pain donors, although there was no difference between pain states in any region (C, ** = p < 0.01, * = p < 0.05). Western blots of GAPDH (D), Homer1 (E), gephyrin (F) and the GABA_A_ α2 subunit (G) revealed no significant differences in the quantity of each protein across the entire dorsal grey and white matter of the spinal cord between non-pain and pain donors (P > 0.05 for all comparisons). NP = non-pain, P = chronic lower limb pain.

While the size of Homer1-IR and gephyrin-IR puncta were consistent between non-pain and pain donors, a change in the quantity of Homer1 or gephyrin proteins could indicate alterations in the density of excitatory or inhibitory synapses, respectively. Protein abundance was therefore quantified by western blot. GAPDH expression did not differ between protein lysates made from the dorsal half of lower lumbar spinal cord sections of the same 10 non-pain and 8 pain donors above (Figure 6D) and was therefore used for normalization.

Protein abundances of Homer1 (Figure 6E) and gephyrin (Figure 6F) were consistent between non-pain and pain donors, suggesting there is no expression change in these proteins. To look closer at inhibitory transmission, we assessed the quantity of the GABA_A_ α2 subunit as this has been a proposed target for the treatment of chronic pain [19; 26; 60], however we found no significant difference in this protein in the dorsal half of the spinal cord between non-pain and pain donors either (Figure 6G). Together, these data provide no evidence for widespread changes in postsynaptic marker size or abundance with pain.

## Discussion

Rodent studies have provided conflicting evidence as to whether a loss of inhibitory interneurons occurs as a result of peripheral nerve injury, which may contribute to the hypersensitivity observed in these models. Shrunken, hyperchromatic inhibitory cells were found in the DH following chronic constriction injury [47], purportedly evidence of degeneration caused by persistent ectopic activity from injured primary afferent fibers [25; 47; 48]. Another group showed reduced inhibitory transmission in the spinal DH, alongside apoptotic cells that were positive for terminal deoxynucleotidyl transferase-mediated biotinylated UTP nick end labeling (TUNEL) staining following peripheral nerve injury [24]. It was subsequently proposed that degeneration of GABAergic neurons was caspase- [40] and glutamate-dependent [17]. However, several stereological studies found no evidence of a loss of neurons or a change in the proportion of excitatory to inhibitory neurons in the spinal DH following peripheral nerve injury, despite animals showing clear signs of neuropathic pain [31–33]. These authors also observed that the TUNEL staining seen after peripheral nerve injury was restricted to microglia [32]. In this study, we have shown that chronic lower limb pain is not associated with a reduction in the number or density of neurons in the human lumbar DH in comparison to donors with no medical history of persistent pain. We have also shown that the ratio of inhibitory to excitatory neurons is consistent throughout the DH between groups. Together, these data support the proposal that there is no specific loss of inhibitory interneurons associated with chronic pain, and therefore other mechanisms aside from inhibitory neuron loss must underly pain sensitization in humans.

Our data show that the cross-sectional area of the human spinal cord and neurons within it do not change with chronic pain, however when normalized to the spinal cord size for each donor, the relative DH area was slightly smaller in those with pain (0.069 ± 0.0069 in non-pain donors, 0.062 ± 0.0049 in donors with pain). There were no individual laminar differences, suggesting there is a minor non-specific reduction across the entire grey matter. The central terminals of some primary afferents have been shown to retract from the dorsal horn in rodent nerve injury models [23; 29]. Interestingly, the lumbar DRGs from the 4 donors with peripheral neuropathy-induced chronic pain all scored high for the presence of Nageotte nodules in a separate study [43]. Therefore, while we detected no loss of neurons in the spinal cord, there is evidence of neuronal degeneration in the DRG of the same donors with peripheral neuropathy. This DRG degeneration could result in a reduction of grey matter volume but evidently does not cause any neuronal degeneration in the DH. Future studies should investigate the organization of central terminals in human spinal cord tissue, as a reorganization at the first synapse in the pain pathway would have strong implications in the establishment of a persistent pain state.

Many studies have investigated spinal cord circuits in the rodent [52] and one of our major aims is to understand if this organization is maintained across species, which we partially addressed in this study. Here, we have shown that inhibitory neurons account for approximately 1/3 of neurons in the human DH, meaning excitatory spinal neurons outnumber inhibitory neurons approximately 2:1, similar to the pattern seen in rodents [30; 33]. These broad groups of DH neurons have been divided into distinct subpopulations based on molecular markers, morphology and function in rodent studies [15; 27; 28; 52]. It is likely that this complexity also applies in the human DH, as several molecular markers, such as *NPY* and *GRPR*, have been shown to be specifically expressed by distinct populations in human spinal cord sequencing studies [13; 59; 61]. Interestingly, we found examples of mixed neurons containing transcripts related to both excitatory and inhibitory neurotransmission in all donors, suggesting some neurons in the human spinal cord are potentially capable of dual transmitter release. These mixed neurons have also been identified in human spinal cord sequencing studies [13; 59; 61] and have been shown to be present throughout the grey matter with spatial transcriptomics [13]. A recent study demonstrated neurons containing both glutamatergic and GABAergic genes are somewhat widespread, being present in more than 30 brain regions in the mouse [58]. Switching of neurotransmitter phenotype has also been proposed to be somewhat plastic such that the main transmitter released from a single neuron can change during development or with injury [54], however, we found that the proportion of the mixed *PAX2*+/*SLC17A6*+ neurons in the DH did not change with chronic lower limb pain. Further research is required to understand the role of these mixed neurons in human sensory transmission.

Projection neurons are another class of spinal neurons that have been intensely studied with rodent projections neurons markers, such as TACR1 and GPR83, showing similar staining patterns in human spinal cord [41; 42]. We used size to provide a threshold for candidate projection neurons as these rare neurons have been shown to have a significantly greater soma cross-section than interneurons [1; 7]. The largest 1% of human DH neurons were restricted to lamina I, the deeper laminae and the white matter in a distribution matching that of the rodent [3; 56]. We have also provided quantitative evidence for the existence of the LSN in human, as all sections from every donor analyzed contained rare NeuN-IR cells in the white matter lateral to lamina II (∼2-3 per 20µm section). Interestingly, we also observed neurons in the white matter in an intermediate zone between the LSN and the RLV region of the spinal cord. The size and packing density of neurons in these regions did not change between groups, therefore more research is required to characterize human projection neurons and assess whether their molecular signature changes with chronic pain.

Our data suggests the size of both excitatory and inhibitory synapses in the human DH does not change with chronic pain, likewise neither does the abundance of key proteins found in the postsynaptic scaffolding mesh. Homer1 is present on both dendritic shafts and dendritic spines [51], the latter of which have been shown to rearrange closer to the neuronal cell body following peripheral or spinal cord injury [49]. Therefore, while the abundance of Homer1 did not change, it is possible that reorganization of Homer1+ puncta toward the soma occurs during chronic pain states in humans, thereby increasing the likelihood of action potential generation from excitatory synaptic input. There may also be differences in the abundance of synaptic proteins between neuronal subpopulations or across laminae, which may not have been detected due to the composition of the protein lysates. To maintain protein integrity, lysates were processed from the entire dorsal half of the spinal cord including both the white and grey matter. Ideally individual laminae would be microdissected to search for more subtle differences, but this would require technical optimization that would sacrifice precious tissue samples. This could also explain why we did not observe a difference in the abundance of the GABA_A_ α2 subunit within the dorsal spinal cord in this study. Synaptosome RNA or protein assessment, or laser capture microdissection followed by proteomics will be required to understand subtle changes in synaptic architecture with pain in humans.

There are some important limitations of our study, including the small sample size. While we did not find any evidence of neuronal loss in the lower lumbar DH in this cohort, it does not rule out the possibility that subtle differences would be observed with a larger sample size, geographically distinct cohorts, or in the sacral segments of the spinal cord, which also receive innervation from the feet. Another limitation is that we have not used ultrastructural techniques to assess synapses or synaptic proteins. This could be an area for fruitful investigation in the future, but our negative results on gross synapse size suggest that any such changes might be at the level of protein arrangement within synapses and not large-scale changes in synaptic architecture.

The difficulty in obtaining high-quality human post-mortem tissue has hindered efforts to unravel the organization of the human nervous system. Even high-quality tissue retrieved within hours of cross-clamp can be technically difficult to work with and some techniques used to study circuitry in the rodent, such as retrograde tracing, are not feasible in humans. Despite the difficulty of using these tissues, the importance of understanding the translatability of rodent research and how human spinal circuitry is impacted by chronic pain cannot be overstated. Pursuant to this goal, we have found no evidence that inhibitory neurons are lost in the spinal cord of patients with chronic lower limb pain, suggesting this is not a key mechanism contributing to prolonged mechanical hypersensitivity and chronic lower limb pain. We also found no evidence that the size and density of key post-synaptic proteins are altered in chronic pain states, suggesting synapses are also stable. Future studies can build on these results to improve our understanding of the long-term changes that occur in the human spinal DH with chronic pain, leading to targeted therapeutic treatments for these debilitating conditions.

## Acknowledgements

The authors thank the organ donors and their families for their gift of life. The authors also thank all members of the UTD hDRG tissue team for help with tissue retrieval. The authors finally thank the previous and current professors of the Spinal Cord Group at the University of Glasgow for access to the frozen bank of antibodies to trial on human spinal cord tissue.

## Data Availability Statement

All data is presented within the paper. Raw image files are available upon request.

## List of supplemental figures

**Supplemental Table 1: List of antibodies trialed to label inhibitory and excitatory neuronal cell bodies.**

**Supplemental Table 2: Summary of laminar distribution and neuronal size across the human dorsal horn**

**Supplemental Table 3: Summary of the proportion of inhibitory and excitatory neurons across the dorsal horn in donors with and without chronic pain**

**Supplemental Table 4: Summary of postsynaptic protein marker analyses across the human spinal dorsal horn**

**Supplemental Figure 1:**
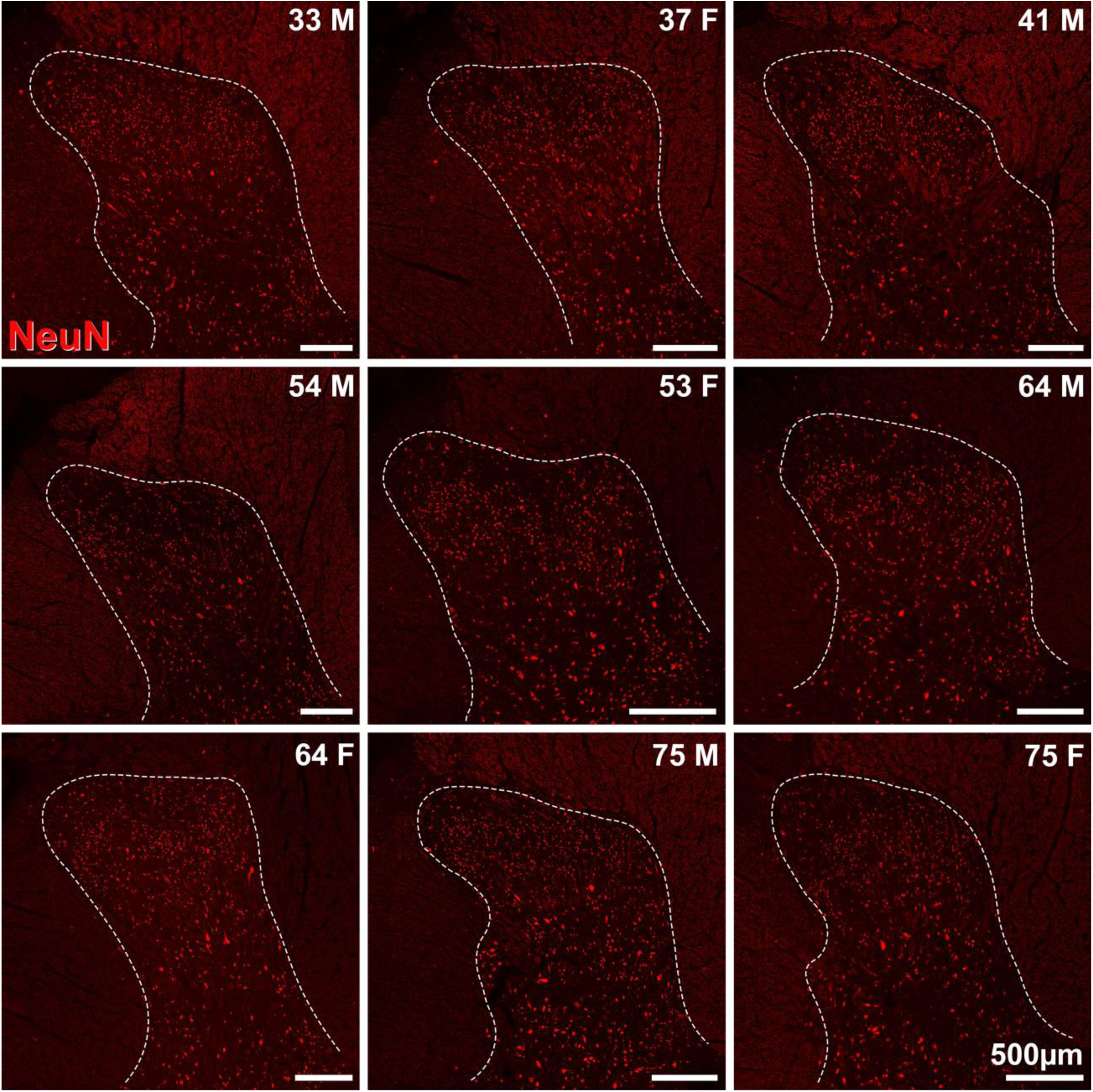
Representative NeuN immunolabelling in the dorsal horn of transverse lower lumbar spinal cord sections from 9 human organ donors with no history of chronic pain. Images are a single optical section, x20 magnification. All individual NeuN+ profiles are hard to detect at this low magnification required to visualize the shape of the dorsal horn, but NeuN immunolabelling from the same tissue sections are shown at a higher zoom in Supplemental Figure 3. F = female; M = male. Numbers equate to age in years.

**Supplemental Figure 2:**
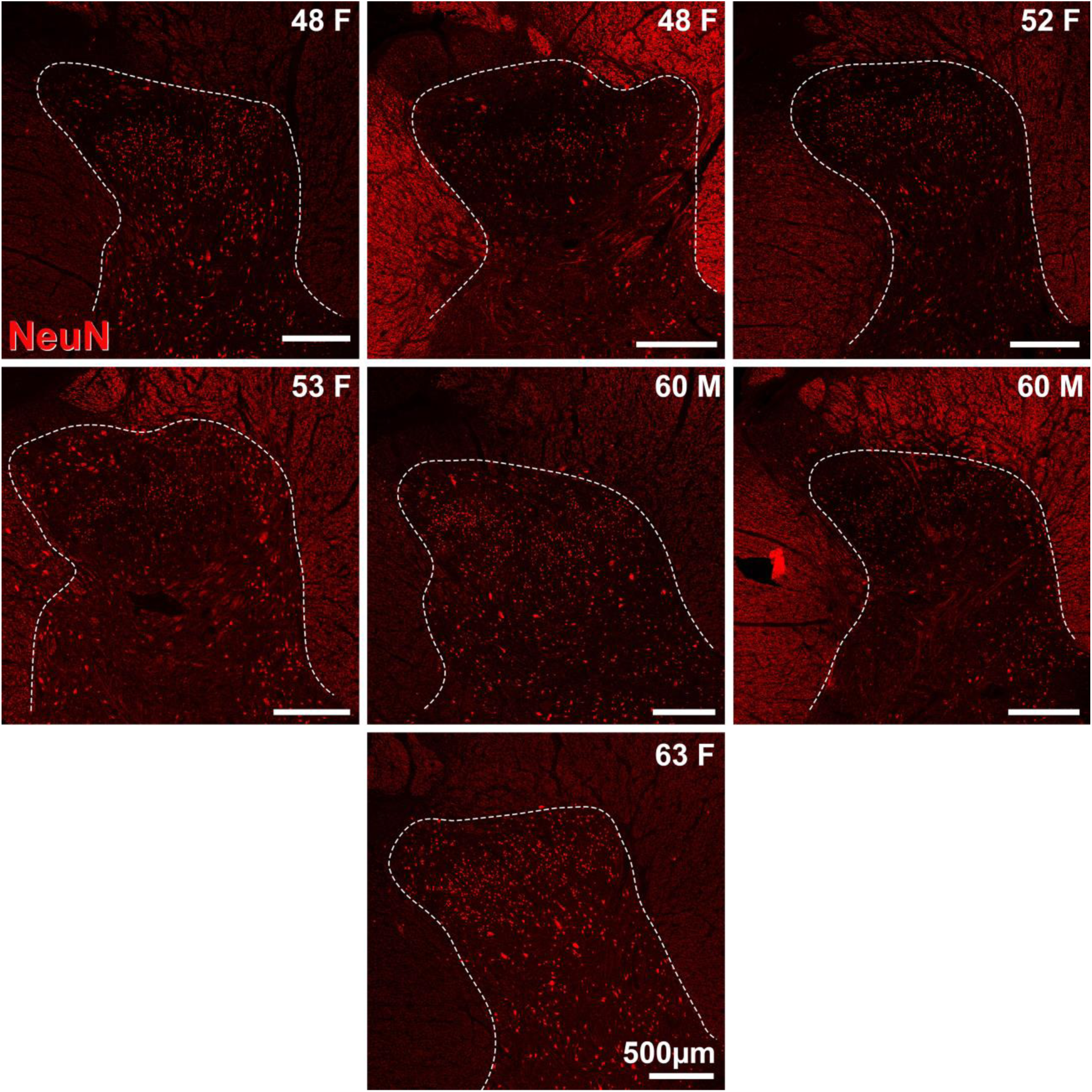
Representative NeuN immunolabelling in the dorsal horn of transverse lower lumbar spinal cord sections from 7 human organ donors with documented lower limb pain. Images are a single optical section, x20 magnification. All individual NeuN+ profiles are hard to detect at this low magnification required to visualize the shape of the dorsal horn, but NeuN immunolabelling from the same tissue sections are shown at a higher zoom in Supplemental Figure 3. F = female; M = male. Numbers equate to age in years. The two 48F and 60M examples are from different individuals.

**Supplemental Figure 3:**
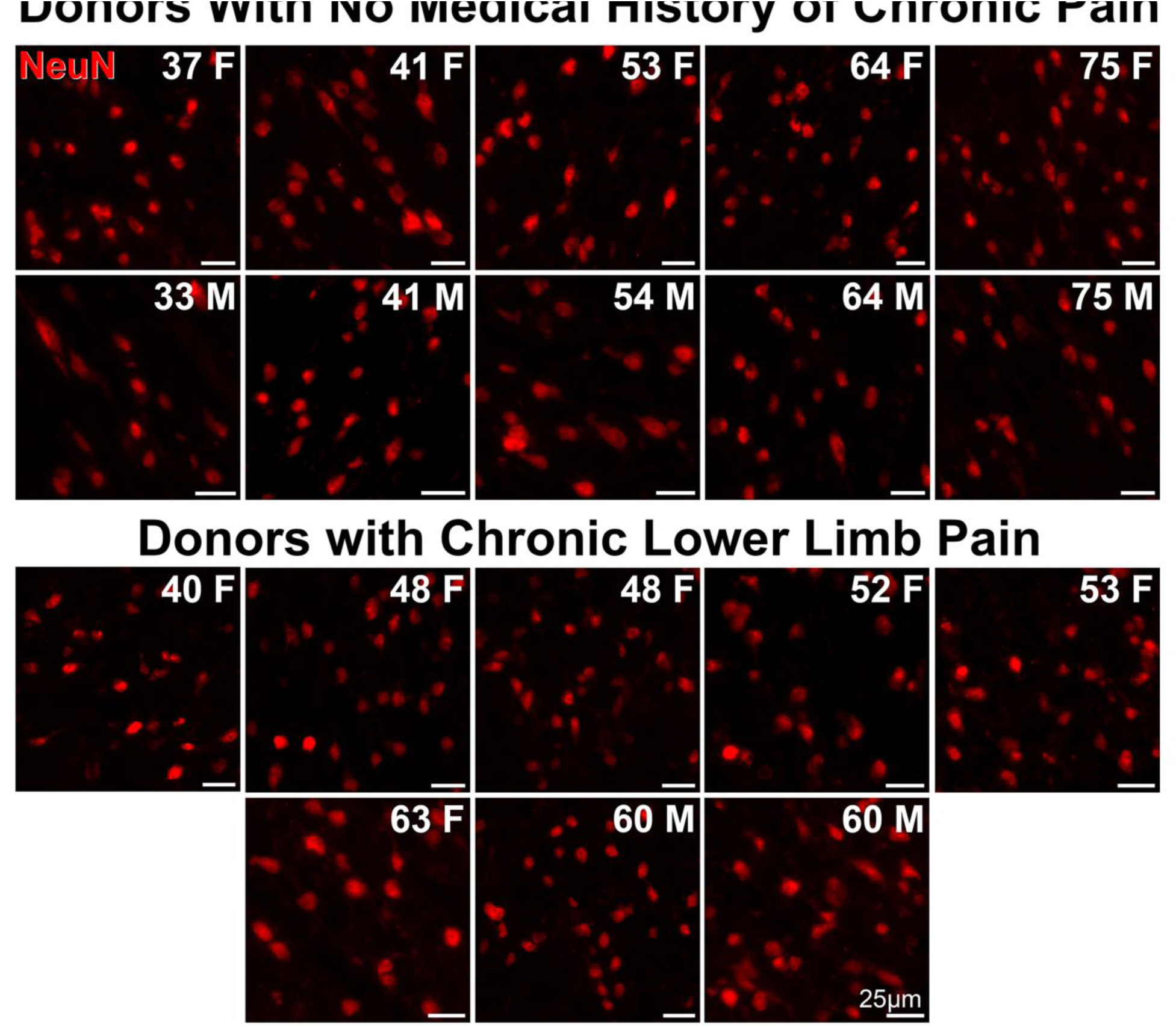
Representative detailed view of NeuN immunolabelling from the superficial laminae of transverse lower lumbar spinal cord sections from 18 human organ donors. Representative images from 10 donors with medical history of chronic pain, and 8 donors with chronic bilateral lower limb pain. Images are cropped from a single optical section of the entire dorsal horn, x20 magnification. F = female; M = male. Numbers equate to age in years. The two 48F and 60M examples are from different individuals.

**Supplemental Figure 4:**
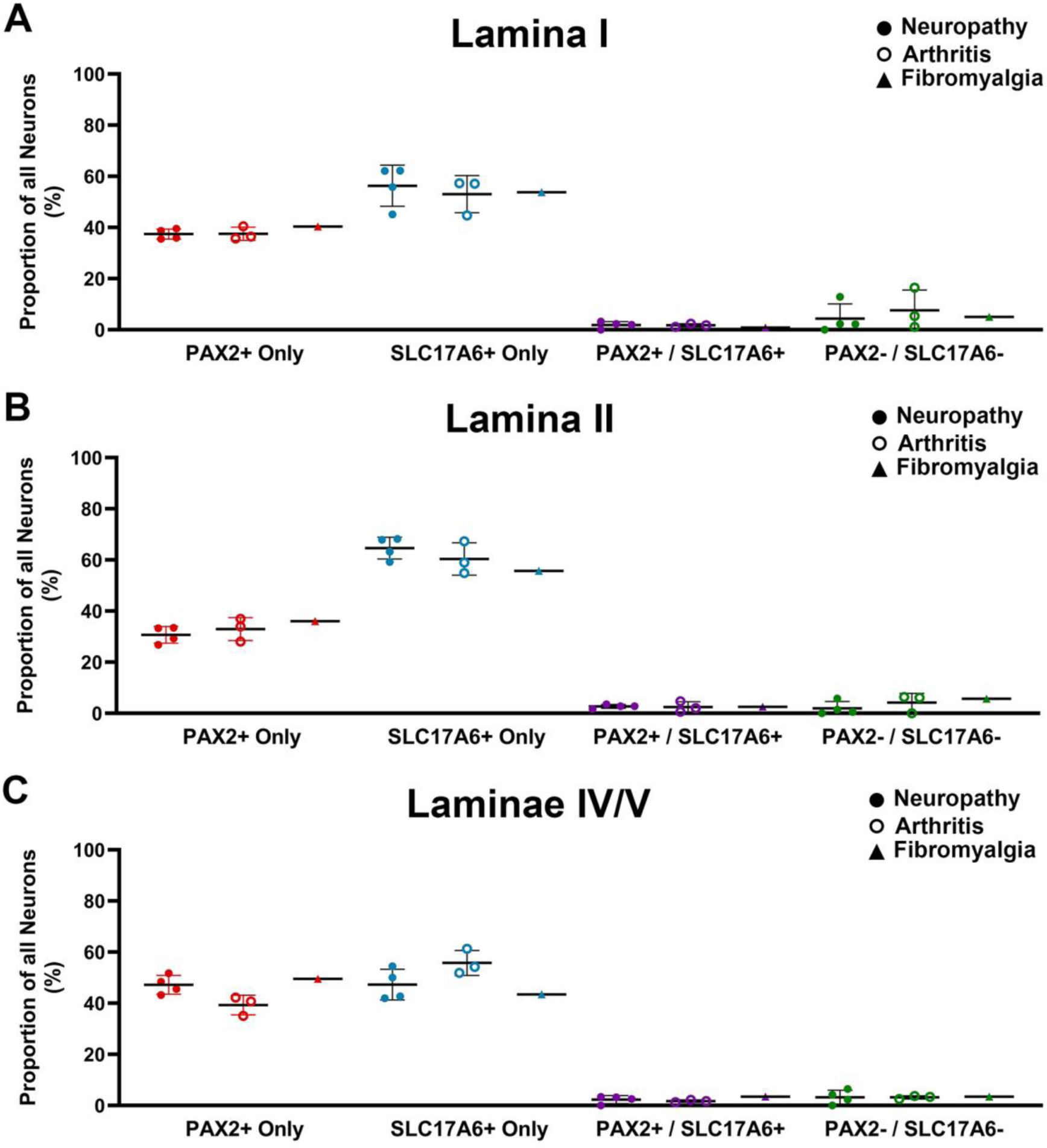
The proportion of excitatory and inhibitory neurons is comparable between donors with neuropathic, arthritic or fibromyalgic derived pain histories. The ratio of *PAX2*+ only, *SLC17A6*+ only, mixed *PAX2*+/*SLC17A6*+ and *PAX2*-/*SLC17A6*-neurons in lamina I (A), lamina II (B) and laminae IV/V (C) is comparable between donors with chronic pain induced by peripheral neuropathy or arthritis conditions.

